# The PCM scaffold enables RNA localization to centrosomes

**DOI:** 10.1101/2024.01.13.575509

**Authors:** Junnan Fang, Weiyi Tian, Melissa A. Quintanilla, Jordan R. Beach, Dorothy A. Lerit

## Abstract

As microtubule-organizing centers, centrosomes direct assembly of the bipolar mitotic spindle required for chromosome segregation and genome stability. Centrosome activity requires the dynamic assembly of pericentriolar material (PCM), the composition and organization of which changes throughout the cell cycle. Recent studies highlight the conserved localization of several mRNAs encoded from centrosome-associated genes enriched at centrosomes, including *Pericentrin-like protein* (*Plp*) mRNA. However, relatively little is known about how RNAs localize to centrosomes and influence centrosome function. Here, we examine mechanisms underlying the subcellular localization of *Plp* mRNA. We find that *Plp* mRNA localization is puromycin-sensitive, and the *Plp* coding sequence is both necessary and sufficient for RNA localization, consistent with a co-translational transport mechanism. We identify regions within the *Plp* coding sequence that regulate *Plp* mRNA localization. Finally, we show that protein-protein interactions critical for elaboration of the PCM scaffold permit RNA localization to centrosomes. Taken together, these findings inform the mechanistic basis of *Plp* mRNA localization and lend insight into the oscillatory enrichment of RNA at centrosomes.

## Introduction

Centrosomes are microtubule-organizing centers (MTOCs) that support cell division, intracellular trafficking, and ciliogenesis. Consequently, centrosome dysfunction is associated with varied diseases and developmental disorders, including cancer and microcephaly [1–3]. Centrosome function is instructed by the organization and composition of the pericentral material (PCM), the composite of proteins and mRNAs that surround the central pair of centrioles [4–7].

Centrosome activity oscillates in phase with the cell cycle. Centrosomes duplicate once and only once per cell cycle, usually during S-phase [8]. Subsequently, the duplicated centrosomes separate and undergo a maturation process, wherein additional PCM is recruited to support microtubule nucleation and organization [9–13]. The coordinated processes of centrosome duplication, separation, and maturation ensure the timely formation of the bipolar mitotic spindle during M-phase. As cells exit mitosis, centrosomes shed PCM [13, 14]. While these cell cycle-dependent fluctuations in PCM recruitment and shedding instruct the microtubule-organizing activity of centrosomes, the underlying mechanisms remain incompletely understood.

Recent work indicates that some mRNAs specifically enrich at centrosomes in a cell cycle-dependent manner [7, 15–18]. Remarkably, RNAs that localize to centrosomes encode centrosome proteins, raising the intriguing possibility that centrosomal mRNAs may contribute to centrosome maturation, structure, or otherwise influence centrosome activity [19–21]. Consistent with these ideas, the localization of some centrosomal mRNAs is directed by a co-translational transport mechanism, whereby RNA localization and protein translation are coupled [15, 22, 16]. Within cultured mammalian cells, for example, *ASPM* and *NUMA1* mRNAs and nascent peptides are co-trafficked to centrosomes followed by additional on-site translation [15]. Co-translational transport was similarly reported for *Centrocortin* (*Cen*) mRNA within *Drosophila* syncytial embryos [22]. Furthermore, the mislocalization of *Cen* mRNA to the anterior cortex prevents Cen protein from localizing to distal centrosomes, demonstrating the coupling of RNA transport and local translation [17].

Among the most conserved mRNAs localizing to centrosomes is *Pericentrin* (*PCNT*) mRNA, as observed in cell culture, zebrafish, and *Drosophila* models [16–18]. Human PCNT and *Drosophila* Pcnt-like protein (Plp) share functionally conserved roles in PCM scaffolding and microtubule nucleation [23–28]. In humans, loss-of-function *PCNT* mutations are associated with microcephalic osteodysplastic primordial dwarfism type II (MOPD II), as well as cardiac and neurovascular abnormalities [29–33]. Loss of *Drosophila Plp* also leads to pleiotropic effects, including embryonic lethality, neuronal dysfunction, and sterility [24, 25, 28, 34]. While prior work indicates *PCNT* mRNA localization requires translation and the microtubule minus end-directed motor dynein, relatively little is understood about mechanisms underlying the co-translational transport of centrosomal RNAs or how their localization is coupled to the cell cycle [16].

*Drosophila* embryos are a valuable model to investigate how and why RNAs localize to centrosomes. *Drosophila* embryos progress through 14 rounds of synchronous, abridged S-to-M nuclear division cycles (NCs) without gap phases prior to somatic cellularization [35]. During this period of development, the embryo is largely transcriptionally quiescent and supported by maternal stores of RNAs and proteins [36]. As in mammalian cells, RNAs enrich at embryonic centrosomes preceding mitotic onset, and less RNA localizes to centrosomes during mitosis [17, 15]. RNAs also progressively enrich at centrosomes as embryonic development ensures, concomitant with the lengthening of successive NCs [17, 18]. These findings argue that RNA localization to centrosomes is entrained to the cell cycle and developmental progression.

Prior work by our group and others similarly uncovered cell cycle and developmental stage-specific changes in the organization of *Drosophila* embryonic centrosomes. The organization and structure of the PCM is largely supported by the formation of Centrosomin (Cnn) flares, which extend during interphase, retract during mitosis, and mature as the NCs proceed [24, 37]. Cnn functions as a PCM scaffold important for centrosome maturation and organization [37–39]. Cnn scaffolding activity, in turn, is supported by Plp, which localizes to the tips of Cnn flares and interacts directly with Cnn via two interaction modules. The interaction between Plp-Cnn is critical for PCM scaffolding and early embryo mitotic divisions [24]. Although the oscillations in centrosomal RNA distributions appear to mirror changes in PCM organization, whether the PCM scaffold influences RNA localization has not been examined.

In this study, we examine the mechanisms that support *Plp* mRNA localization to centrosomes. We show *Plp* mRNA localization is puromycin-sensitive, consistent with a co-translational transport mechanism. We further identify a requirement for microtubules to direct *Plp* mRNA to centrosomes. Through a reporter assay, we discovered the *Plp* untranslated regions are dispensable for *Plp* mRNA localization. Rather, regions within the *Plp* coding sequence (CDS) necessary for PCM scaffolding also direct mRNA localization. We further demonstrate genetic perturbation of the PCM scaffold is sufficient to impair centrosomal mRNA localization. Taken together, these data inform mechanisms underlying *Plp* mRNA localization and the basis of cell cycle-dependent variances in RNA enrichment at centrosomes.

## Results

### Microtubules support *Plp* mRNA localization

Centrosomes are MTOCs, and RNA localization often utilizes microtubule-based transport, raising the possibility that microtubules help enrich RNA at centrosomes [40, 41]. To investigate the role of microtubules in the subcellular localization of *Plp* mRNA at centrosomes, we performed microtubule regrowth assays [42]. Microtubule stability is sensitive to temperature; therefore, microtubules were depolymerized by incubating early embryos on ice (see Methods) [43–45]. We first confirmed that cold-shock treatment led to microtubule depolymerization and the loss of Cnn flares, consistent with prior work [24, 46]. To allow microtubule regrowth, we shifted cold-shocked embryos to room-temperature, which also permitted reformation of Cnn flares (**Figure 1A**).

**Figure 1.**
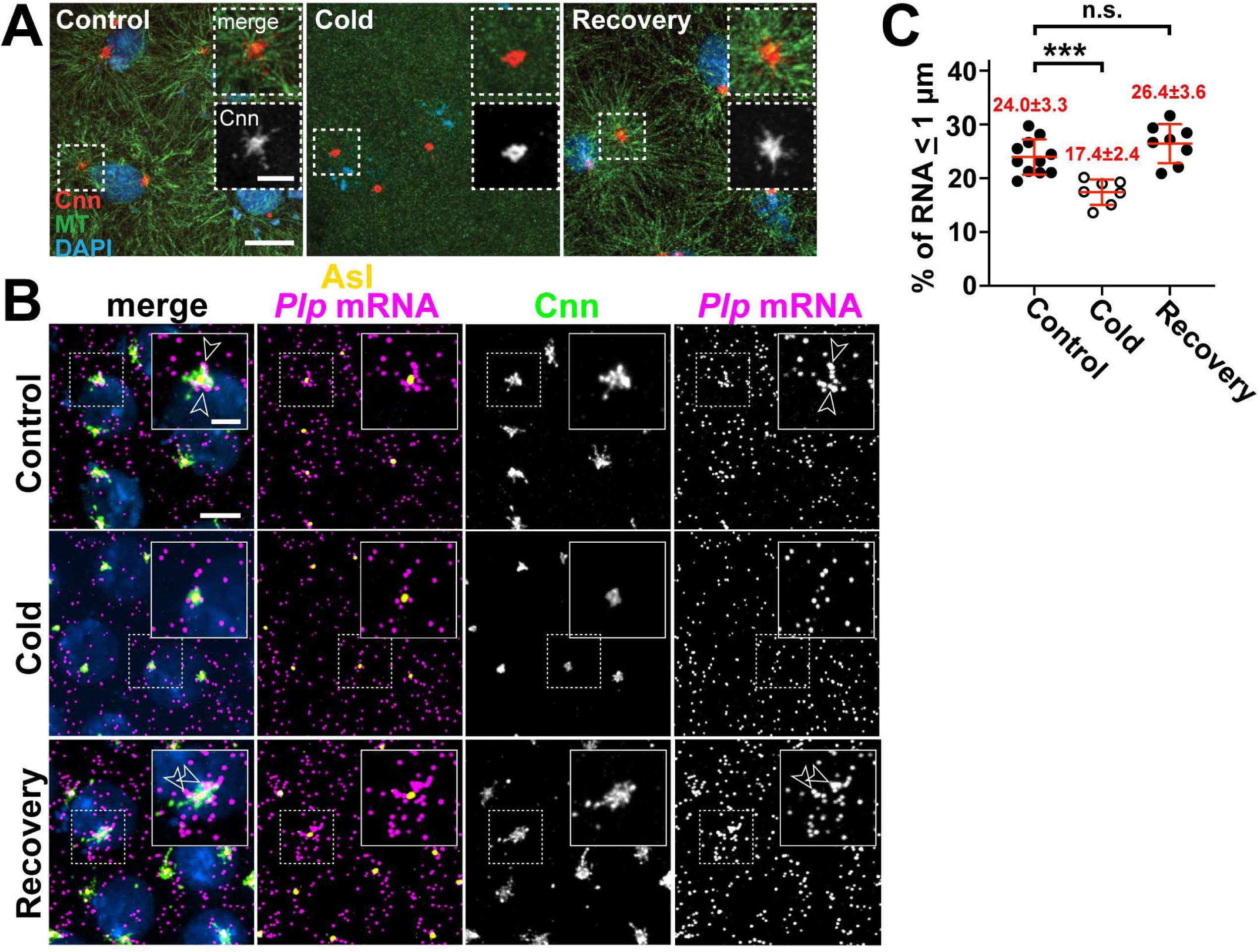
*Pip* mRNA localization requires microtubules. Maximum intensity projections of (A) NC 11 embryos from the indicated conditions labeled with anti-Cnn (PCM; red) and a-Tub antibodies (microtubules; green), and DAPI (DNA; blue). (B) NC 12 *Plp-GFP* embryos from control, cold-treated, or recovery conditions labeled with *GFP* smFISH probes to show *Pip* mRNA distributions (magenta) and labelled with Cnn (green) and Asl (centrioles; yellow) antibodies and DAPI (blue). Dashed box regions are enlarged in insets. Arrowheads show *Pip* mRNA enrichments at centrosomes. (C) Quantification of *GFP* mRNA localization (within 1 µm from Asl). Each dot represents a measurement from a single embryo; see Table S1 for N embryos and RNA objects examined. Mean ± S.D. are displayed. n.s. not significant; ***p<0.001 by one-way ANOVA followed by Dunnett’s T3 multiple comparisons test. Scale bars: 5 µm; 2 µm (insets).

Microtubule depolymerization decreased endogenous *Plp* mRNA localization, as revealed by single molecule fluorescence in situ hybridization (smFISH). This response was reversible, as *Plp* mRNA localization was restored to WT levels following microtubule regrowth (**Figure 1B,C**). Thus, microtubules support proper *Plp* mRNA localization to centrosomes.

Cytoplasmic dynein is a minus-end directed microtubule motor, reviewed in [47]. Prior work established a requirement for dynein to localize *PCNT* mRNA and protein to centrosomes in cultured human cells [16, 48, 49]. PCNT associates with dynein via a dynein light intermediate chain (DLIC) recognition motif situated in the middle of the *PCNT* CDS [50]. By sequence analysis, we confirmed that this region contains an AAxxG motif important for DLIC recognition [51]. In contrast, *Drosophila* Plp and mouse Pcnt lack the AAxxG motif, indicating this region is less well conserved (**Figure S1A**).

To directly test whether dynein similarly functions in translocating *Plp* mRNA to centrosomes, we examined RNA distributions in hypomorphic *Dynein heavy chain 64C* (*Dhc*) mutant embryos (i.e., *Dhc^LOA^* homozygous mutants; see Methods). Unexpectedly, we did not observe significant changes to *Plp* mRNA localization in *Dhc^LOA^* mutants relative to controls (**Figure S1B, C**). These findings suggest that either sufficient dynein activity persists in *Dhc^LOA^*mutants or that other mechanisms support *Plp* mRNA localization.

### Co-translational transport of *Plp* mRNA to centrosomes

We previously showed some *Plp* mRNA colocalizes with Plp protein at centrosomes [18]. Consistent with these observations, recent work highlights co-translational transport as a major paradigm for RNA localization to centrosomes [16, 15, 20]. To assess whether translation is required for *Plp* mRNA localization, we examined *Plp* distributions following treatment with several translation inhibitors [52].

Puromycin is an A-site tRNA analog that terminates translation and induces ribosome dissociation from the nascent polypeptide [53]. In contrast, anisomycin and cycloheximide (CHX) block translation elongation and freeze ribosomes on mRNAs without releasing the newly synthesized peptide [54–56]. Treatment with these inhibitors revealed *Plp* mRNA localization is selectively puromycin-sensitive (**Figure 2A, B**). These results argue that actively engaged ribosomes in association with the nascent peptide are drivers of *Plp* mRNA localization to centrosomes, similar to human *PCNT* mRNA [16].

**Figure 2.**
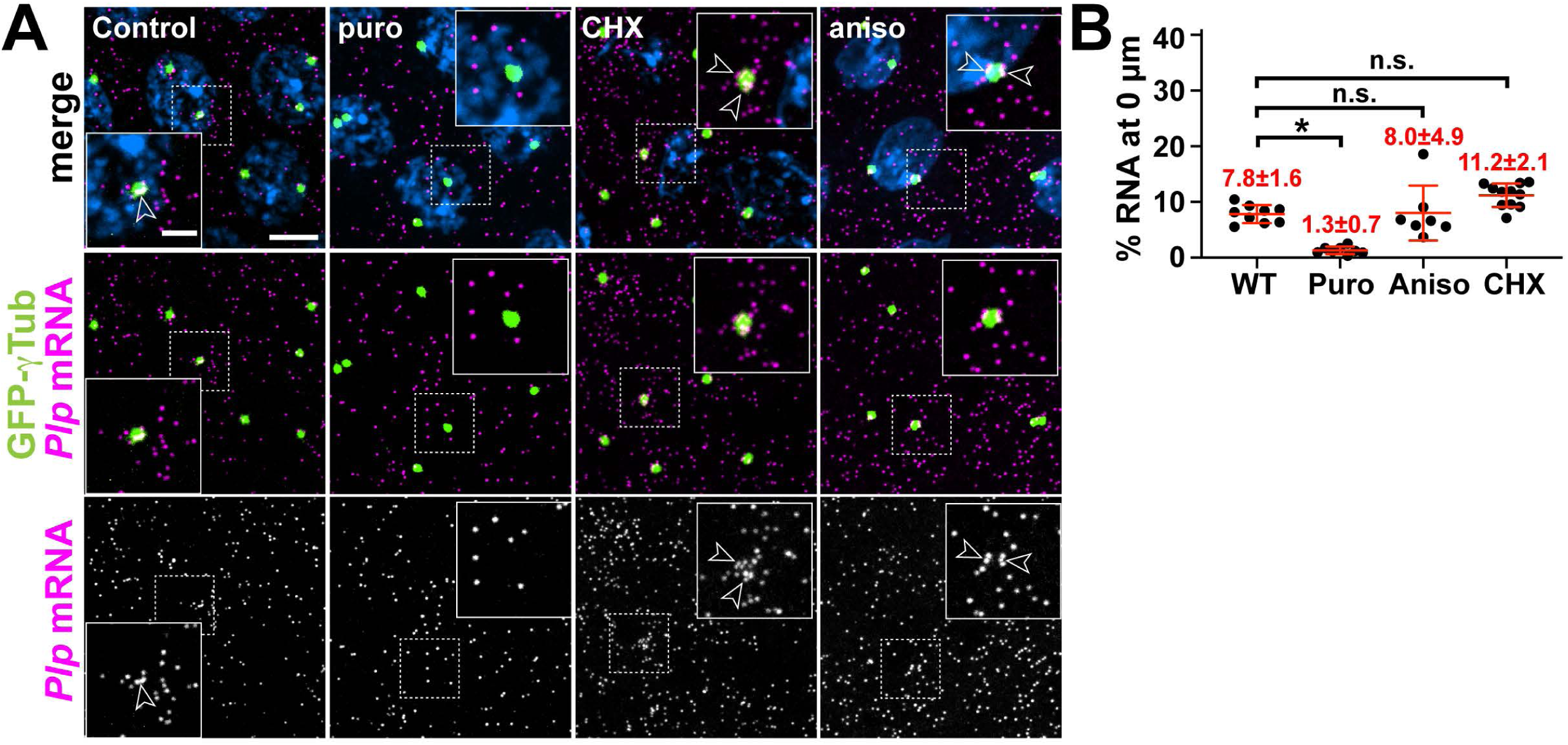
*Pip* mRNA localization to centrosomes is puromycin-sensitive. (A) Maximum intensity projections of NC 13 embryos expressing *GFP-yTub* (green) labeled with *Pip* smFISH probes (magenta) and DAPI (blue) in controls or following treatment with translation inhibitors: puromycin (puro), cycloheximide (CHX), or anisomycin (aniso). Dashed box regions mark insets. Arrowheads show *Pip* mRNA enrichments at centrosomes. (B) Percentage of *Pip* mRNA localizing within O µm from the *yTub* surface. Mean± S.D. are displayed. n.s. not significant; *p<0.05 by one-way ANOVA followed by Dunnett’s T3 multiple comparisons test. Scale bars: 5 µm; 2 µm (insets).

### Domains within the *Plp* CDS direct *Plp* mRNA localization

To further investigate the molecular mechanisms of *Plp* mRNA localization, we utilized a reporter assay to define *cis*-regulatory elements. As a control, we first examined the localization of endogenous *Plp-GFP*, an in-frame GFP knock-in at the Plp C-terminus generated via CRISPR (hereafter, *Plp-GFP*), as schematized in **Figure S2A** [57]. RNA distributions for *Plp-GFP* resembled those of untagged *Plp* mRNA, confirming that the addition of the GFP tag did not alter RNA localization or expression (**Figure S2B,C**). We then used the *maternal α-Tub* driver (*matGAL4*) to direct expression of various *GFP* reporter transgenes and visualized RNA distributions. Because RNA localization often relies upon sequences and/or structural motifs within the untranslated regions (UTRs) [40], we first examined whether the *Plp* 5’- and/or 3’-UTRs were sufficient to localize *GFP* mRNA to centrosomes. Neither the *Plp* 5’- nor 3’-UTRs directed RNA localization to centrosomes, despite expression levels comparable to controls, suggesting that the localization elements reside within the *Plp* CDS (**Figure S2D, E**).

Aligned with this prediction, the *Plp* CDS was sufficient for RNA localization to centrosomes (**Figure 3A,B**; *Plp^FL^-GFP*). This enrichment was specific and not due to spurious overlap because it was eliminated by rotating the RNA channel by 90° (**Figure 3B**). Comparing RNA distributions in *Plp-GFP* versus *Plp^FL^-GFP* embryos indicates the *Plp* CDS mediates localization less efficiently, suggesting that while the *Plp* CDS encodes sequences necessary and sufficient for RNA localization to centrosomes, other features (e.g., regulatory sequences, splicing events, etc.) influence the extent of RNA enrichment (**Figure 3A,B and Figure S2E).** Nevertheless, a requirement for the *Plp* CDS for RNA localization is consistent with the puromycin-sensitivity noted above.

**Figure 3.**
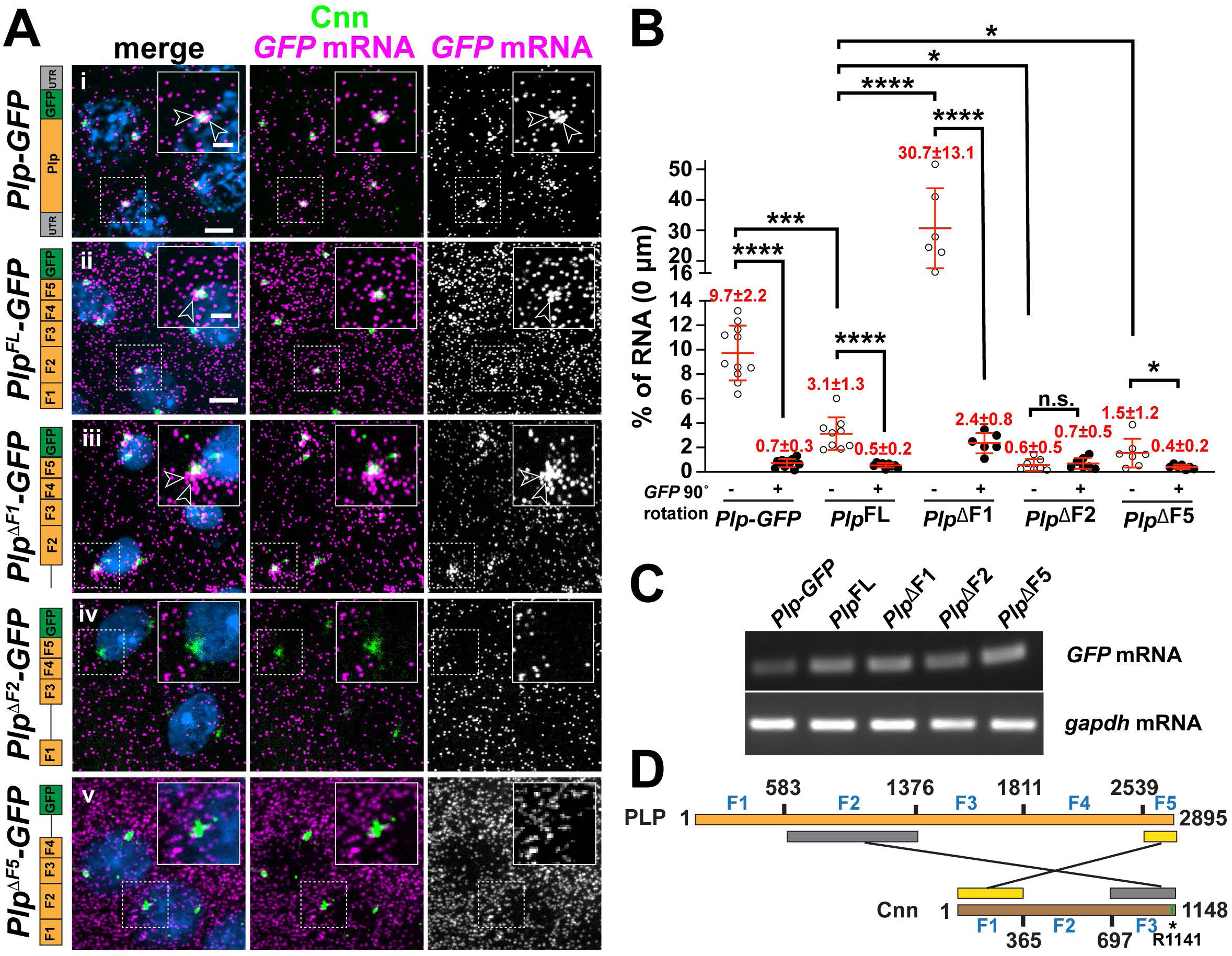
*Pip* mRNA localization requires sequences within the *Pip* CDS. (A) Maximum intensity projections of NC 11 embryos expressing *mCherry-Cnn* (green) and labeled with *GFP* smFISH probes (magenta) to mark transgenic *Pip* mRNA localization and DAPI (blue) in the following genotypes: (i) *Plp-GFP,* (ii) *UAS-PlpFL-GFP,* (iii) *UAS-Plp!iF1-GFP,* (iv) *UAS-Plp!iF2-GFP,* and (v) *UAS-Plp!iF5-GFP.* Transgenes in (ii-v) were expressed using *matGAL4* in the presence of endogenous *Pip.* Construct schematics are shown to the left. Arrowheads show RNA enrichments at centrosomes. (B) Quantification of *GFP* mRNA localization (0 µm from Cnn surface). Each dot represents a measurement from a single embryo; see Table S1 for N embryos and RNA objects examined. The RNA channel was rotated 90° (+) and images re-quantified to assay the specificity of localization. (C) RT-PCR was used to assay the relative expression of the indicated GFP-tagged constructs from 0-2 hr embryos. (D) Schematic adapted from (Lerit et al., 2015) showing the two direct interaction modules between Pip and Cnn. Asterisk denotes the single point mutation (R1141H) that defines the *cnn^84^* allele and abolishes the direct interaction between Pip F2 and Cnn CM2 (green bar). Mean ± S.D. are displayed. n.s. not significant; *p<0.05; ***p<0.001; ****p<0.0001 by one-way ANOVA followed by Dunnett’s T3 multiple comparisons test. Uncropped gels are available at <10.6084/m9.figshare.24926298>. Scale bars: 5 µm; 2 µm (insets).

To uncover which regions of the *Plp* CDS direct RNA localization, we leveraged several *Plp* truncation lines, which divide the ORF into five fragments (F1–F5) and incorporated them into our reporter assay [24, 58] (**Figure 3**). The truncation lines were all overexpressed relative to *Plp-*GFP, but comparable to *Plp^FL^-GFP* (**Figure 3C**). Unexpectedly, we found that an N’-terminal truncation of *Plp* (*Plp*^Δ*F1*^*-GFP*) resulted in significantly more *Plp* mRNA at centrosomes, suggesting that elements within F1 somehow limit *Plp* mRNA localization to centrosomes. In contrast, deletion of either F2 (*Plp*^Δ*F2*^*-GFP*) or F5 (*Plp*^Δ*F5*^*-GFP*) resulted in significantly less *Plp* localized to centrosomes (**Figure 3A,B**). Taken together, these results suggest that expression levels alone are insufficient to instruct RNA localization to centrosomes. Rather, RNA localization to centrosomes is driven by discrete *cis*-elements. In particular, *Plp*^Δ*F2*^ abolished *Plp* mRNA localization, indicating that F2 is required for *Plp* mRNA localization or anchorage at centrosomes.

### The PCM scaffold anchors RNAs at centrosomes

We previously reported that Plp F2 and F5 mediate direct protein-protein interactions with Cnn F3 and Cnn F1, respectively, to maintain the PCM scaffold [24]. The PCM scaffold is impaired in *cnn^B4^* mutants, defined by an R1141H substitution within the highly conserved Cnn Motif 2 (CM2) and sufficient to block the interaction between Plp F2 and Cnn F3 (**Figure 3D**) [59, 24]. Using super-resolution microscopy, we found that *Plp* mRNA appeared displaced from the fragmented PCM in *cnn^B4^*mutants, as compared to age-matched controls (**Figure 4A**). Quantification revealed a significant reduction in *Plp* mRNA localizing within 1 µm from the centriole (marked with Asterless, Asl) in NC 13 *cnn^B4^*mutants, as compared to controls (22.8±8.1% in WT vs. 8.6±4.7% in *cnn^B4^*; **Figure 4B,C**). A similar reduction was observed in early NCs (**Figure S3 A,B)**. Because total levels of *Plp* mRNA are similar in 0–2 hr WT and *cnn^B4^* embryos (**Figure 4D**), we conclude that the PCM scaffold is required to anchor *Plp* mRNA at centrosomes, likely via protein-protein interactions between Plp and Cnn.

**Figure 4.**
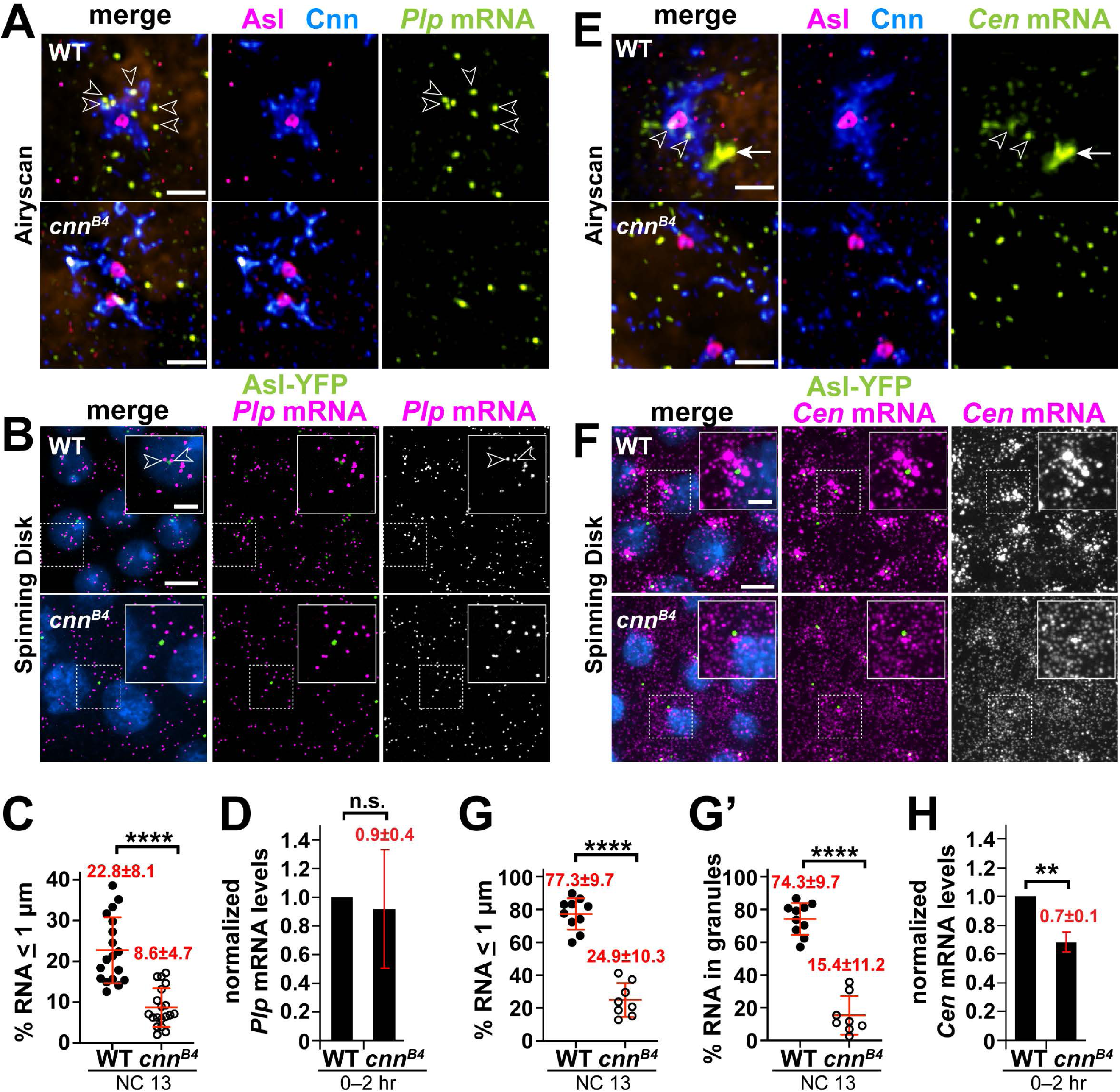
The centrosome scaffold permits mRNA localization. Maximum intensity projections of NC 13 control and *cnn^84^* embryos labeled with (A and B) *Pip* mRNA or (E and F) *Gen* mRNA smFISH probes. In (A and E), embryos were co-stained with smFISH probes (green), anti-Cnn (blue) and Asl (magenta) antibodies, and DAPI (orange; nuclei), then imaged using a Zeiss LSM 880 Airyscan. For (B and F), embryos expressing *Asl-YFP* (green) were labeled with smFISH probes (magenta) and DAPI (blue) then imaged by spinning disk confocal microscopy. (C) Quantification shows the percentage of *Pip* mRNA localizing within 1 µm from the Asl surface. (D) Levels of *Pip* mRNA or (H) *Gen* mRNA were normalized to *RP49* as detected by qPCR from 0-2 hr WT versus *cnn^84^* embryos and displayed relative to the WT control. (G) The percentage of *Gen* mRNA localizing and (G’) residing within granules (defined as 4 RNA molecules per object [17]) within 1 µm from the Asl surface. Each dot represents a measurement from a single embryo; see Table S1 for N embryos and RNA objects examined. Mean± S.D. are displayed. n.s., not significant or ****p<0.0001 by

Might the PCM scaffold support the localization of other centrosome-localized RNAs? Normally, *Cen* mRNA becomes significantly enriched at interphase NC 13 centrosomes within micron-scale granules. However, *Cen* mRNA granules appear diminished in *cnn^B4^* mutants [17]. Indeed, super-resolution imaging revealed fewer and smaller *Cen* mRNA granules in *cnn^B4^* embryos, as compared to controls (**Figure 4E**). Quantitative analysis confirmed significantly less *Cen* mRNA resides within granules or localizes to centrosomes in *cnn^B4^* versus controls (**Figure 4F–G’; Figure S3C–D’**).

We next examined whether this reduction in *Cen* mRNA localization might be attributed to changes in RNA abundance by qPCR. While *Cen* RNA levels are about 30% reduced in 0–2 hr embryonic extracts from *cnn^B4^* mutants relative to WT, this difference is unlikely to account for the 3-fold reduction in RNA localization to centrosomes (**Figure 4G,H**). Taken together, these data suggest that an intact PCM scaffold also contributes to *Cen* mRNA localization, perhaps by stabilizing *Cen* RNA granules. Future work is required to investigate the relationship between Cnn and *Cen* RNA granule formation and whether the *Cen* granule regulates *Cen* mRNA stability.

As a whole, these studies help establish a generalizable requirement for the PCM scaffold to dock localized RNAs at the centrosome. We sought to further test this model using an independent approach. Plp functionally cooperates with Cnn to ensure PCM scaffolding. Thus, loss of *Plp* also leads to a PCM fragmentation phenotype [24, 25]. We therefore examined *Cen* mRNA localization within *Plp* null embryos derived from germline clones (*Plp^GLC^* embryos; see Methods). The significant reduction in *Cen* mRNA localization to centrosomes and residence within granules in *Plp^GLC^*embryos relative to controls support a model wherein the PCM scaffold functions not only in the organization of PCM proteins, but for localized mRNAs as well (**Figure 5A–C**).

**Figure 5.**
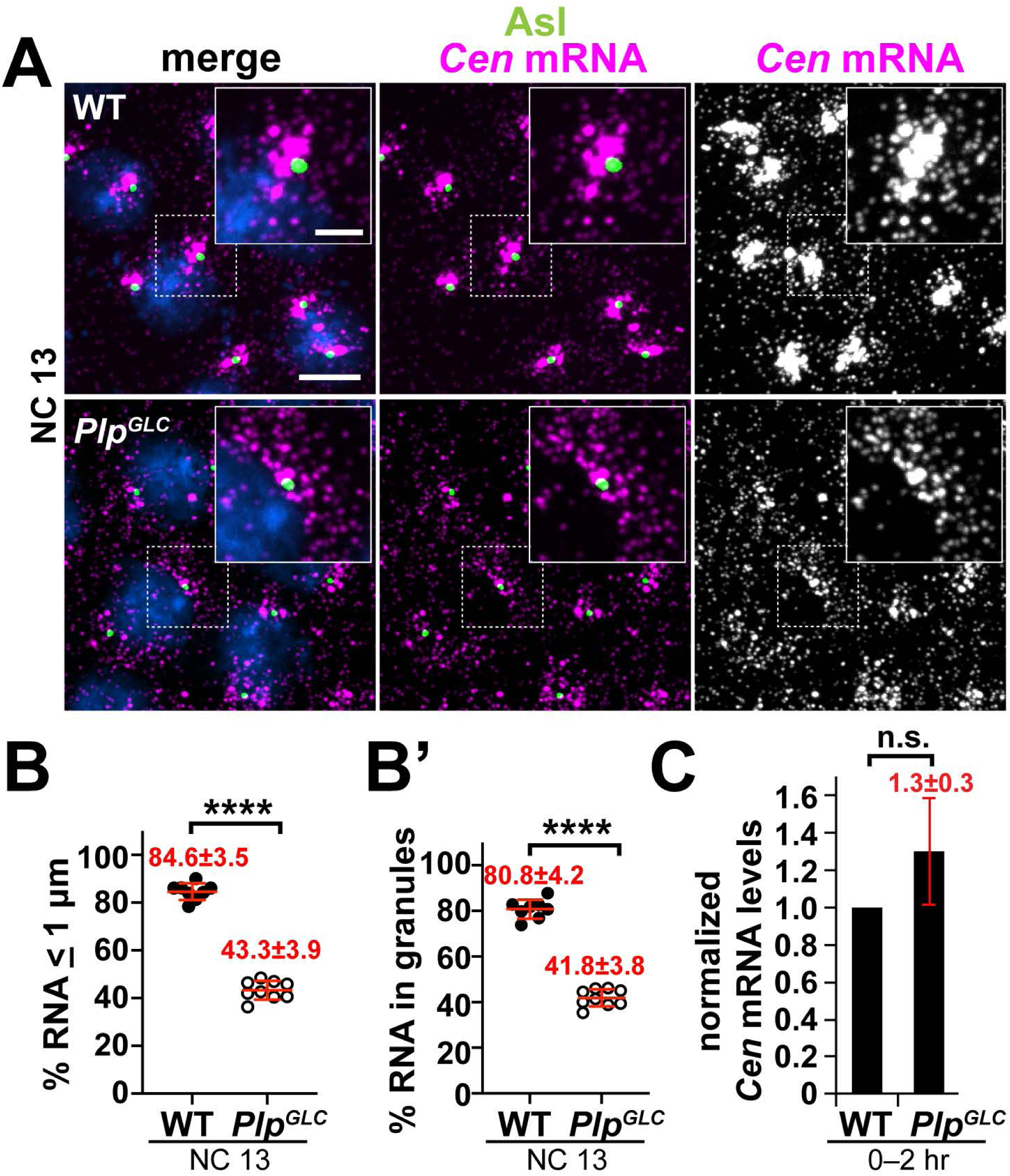
The centrosome scaffold supports *Cen* **mRNA** localization and granule formation. Maximum intensity projections of NC 13 (A) WT or (8) *PfpGLc* embryos labeled with *Cen* smFISH probes (magenta), Asl antibodies (green), and DAPI (blue). Charts show the percentage of *Gen* mRNA (8) localizing or (8’) residing within granules (;:: 4 RNA molecules per object) within 1 µm from the Asl surface. Each dot represents a measurement from a single embryo; see Table S1 for N embryos and RNA objects examined. (C) Levels of *Gen* mRNA were normalized to *RP49* mRNA as detected by qPCR from 0-2 hr WT versus *PfpGLc* embryos and displayed relative to the WT control. Mean± S.D. are displayed. n.s. not significant, and **MTs**

## Discussion

Although some RNAs localize to centrosomes, relatively little is known about their mechanism of localization and function. In this study, we examined the mechanisms of *Plp* mRNA localization to centrosomes. We found that *Plp* mRNA localization requires microtubules, association with the nascent peptide, and defined permissive and restrictive localization elements within the *Plp* CDS. Our findings are consistent with the idea that *Plp* mRNA localization is supported by a protein-protein interaction between Plp F2 and Cnn CM2. We propose that emergence of Plp F2 from the ribosome renders the *Plp* mRNA-protein complex sufficient to associate with Cnn (**Figure 6**), effectively recruiting *Plp* mRNA to centrosomes. Finally, we demonstrated a general requirement for the PCM scaffold to support RNA localization at centrosomes.

**Figure 6.**
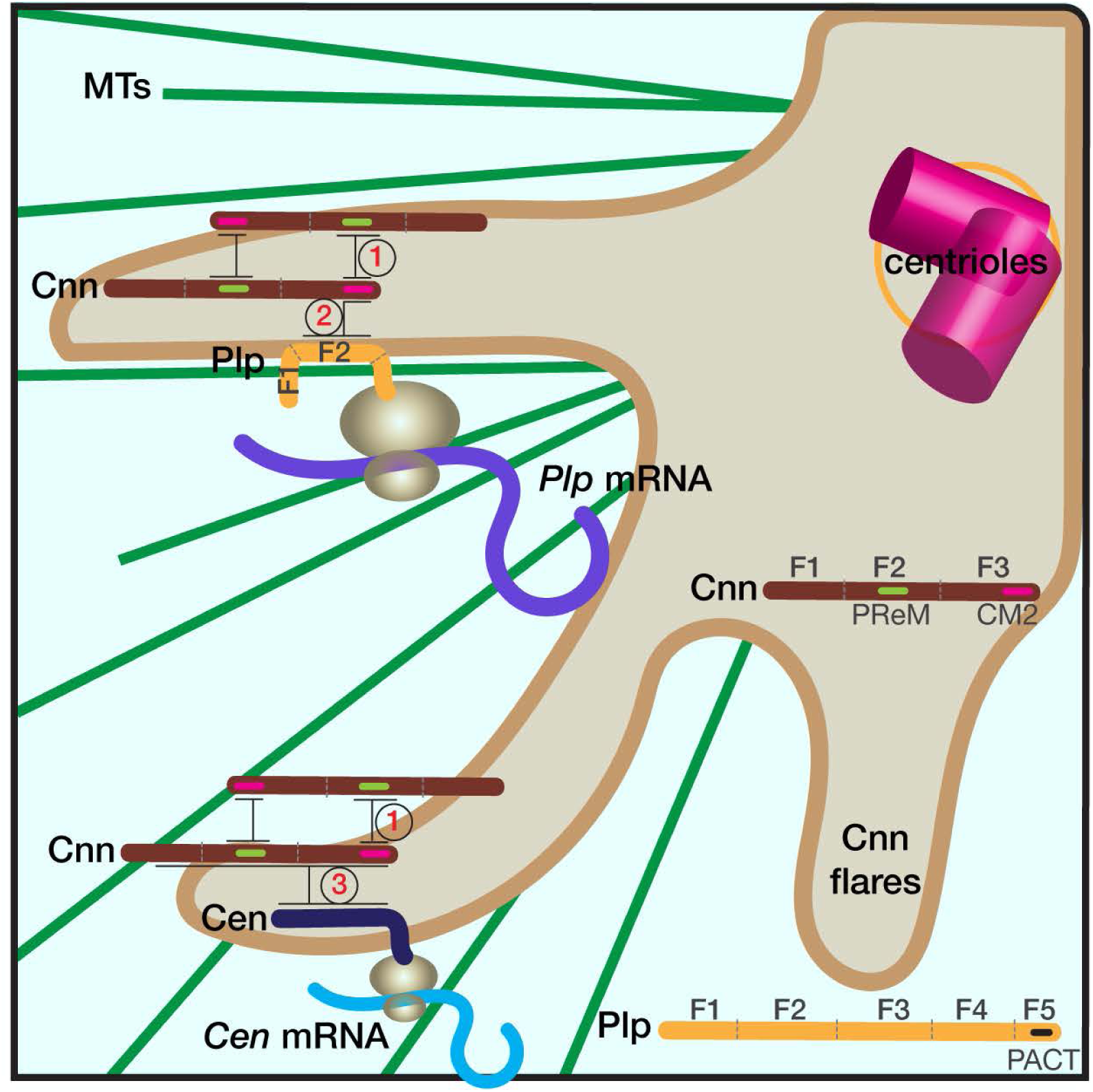
Schematic of RNA localization to centrosomes. The cartoon shows part of a centrosome with extended Cnn flares (brown), which contribute to PCM scaffolding. Elaboration of the PCM scaffold requires oligomerization of Cnn between its PReM and CM2 motifs (interaction 1), and a direct interaction between CM2 and PLP F2 (interaction 2; (61, 39, 24]). Simplified protein architectures of Cnn and Pip are noted in the figure. We propose that the Pip F2-Cnn CM2 interaction helps transmit and/or anchor the *Pip* mRNA-protein complex to the centrosome. Accordingly, microtubules (MTs, green), are required both for the extension of Cnn flares and the localization of *Pip* mRNA to centrosomes (24, 46, 39, 62]. *Cen* mRNA also localizes to the centrosome via co­ translational transport, and Cen protein interacts directly with Cnn (interaction 3). Mutant analysis indicates that an intact PCM scaffold is required for the localization of both *Pip* and *Cen* mRNAs. We further propose that the temporal control of PCM scaffold elaboration (i.e., extension of Cnn flares) similarly regulates RNA localization to centrosomes.

Surprisingly, we found an N’-terminal deletion of Plp F1 led to a significant increase in *Plp* mRNA localization. Recent work demonstrates that the F1 deletion stabilizes Plp, leading to increased protein levels, raising the possibility that the upregulated Plp protein levels in *Plp*^Δ*F1*^ mutants might drive *Plp* mRNA enrichment at centrosomes [60]. However, deletion of F2 led to a significant reduction in *Plp* mRNA localization to centrosomes, despite a similar uptick in Plp protein levels [60]. These findings argue that protein expression levels alone do not direct RNA enrichment at centrosomes. It is more likely that a specific element in F1 limits *Plp* mRNA localization. Future investigation will uncover how Plp F1 suppresses the recruitment of *Plp* mRNA to centrosomes.

In contrast, we found Plp F2 is necessary for *Plp* mRNA localization. This observation is intriguing given our prior work indicating a direct interaction between Plp F2 and Cnn F3, via the CM2, supports centrosome scaffolding and mitotic fidelity [24]. Cnn CM2 promotes centrosome scaffolding through its interaction with a leucine zipper region within a previously identified phosphoregulated-multimerization (PReM) domain residing in the middle of the Cnn CDS (**Figure 6**, *interaction 1*). Phosphorylation of the PReM domain by polo kinase promotes interaction with Cnn CM2, contributing to Cnn oligomerization and scaffold formation [39, 61]. This phosphoregulation likely regulates the timing of centrosome scaffold assembly. Our data indicate *Plp* mRNA localization requires the Cnn scaffold, suggesting the cell cycle-dependent enrichments of *Plp* mRNA at centrosomes are likely entrained to centrosome scaffold formation (**Figure 6**, *interaction 2*).

We also uncovered a requirement for microtubules to support *Plp* mRNA localization. Of note, extension of the centrosome scaffold is also microtubule-dependent, as microtubule depolymerization results in the retraction and condensation of Cnn flares (**Figure 1A**) [24, 46, 39, 62]. In principle, microtubules may be required for *Plp* mRNA localization because they are necessary for scaffold formation. Alternatively, microtubules may help traffic and/or anchor *Plp* mRNA to centrosomes. Live imaging the dynamics of *Plp* mRNA will help decipher these requirements.

Which feature(s) within Plp F2 mediate *Plp* mRNA localization await identification. The recent development of AlphaFold2 allows us to render predictive models of the Plp F2–Cnn CM2 interface. The CM2 within Cnn F3 is critical for centrosome scaffold formation and the interaction with Plp F2, which can be abolished by the *cnn^B4^* R1141H mutation [24, 59, 61, 39]. Using AlphaFold Multimer, an extension of AlphaFold2, and the COSMIC2 cloud platform, we modeled the interface between Plp F2 and Cnn CM2, which provided five predictive structural models [63, 64]. Cnn exists as a monomer in the cytoplasm [65]. Underscoring the fidelity of the AlphaFold predictions, our Cnn CM2 models are similar to the previously reported 3D crystal structure of the CM2 monomer (PDB: 5MVW), with a root mean square deviation (RMSD) ranging from 0.8 to 1.4, confirming the two superimposed atomic coordinates are similar (**Figure S3E**) [61]. We centered our analysis of the AlphaFold models on the Plp F2 residues proximal to Cnn CM2. Intriguingly, all models predicted a C-terminal region of Plp F2 (amino acids 1177– 1306) apposed Cnn CM2 (**Figure S3F**). We speculate that this region is important for the Plp F2 and Cnn F3 interaction and key for *Plp* mRNA localization. While these predictions suggest that Cnn interacts with Plp as a monomer, this requires experimental validation.

Another interacting partner of Cnn is Cen, although the precise interaction interface remains less defined (**Figure 6**, *interaction 3*) [59]. We speculate that this protein-protein interaction similarly supports *Cen* mRNA localization to centrosomes. Given *Cen* and *Plp* mRNA localization both require an intact centrosome scaffold, RNA enrichments are probably temporally coordinated with PCM organization (e.g., entrained to elaboration of the Cnn-rich PCM flares). Nonetheless, the distributions of *Cen* and *Plp* mRNAs are distinct. *Cen* mRNA organizes within large RNA granules, whereas *Plp* mRNA does not. In addition, the localization of the *Cen* mRNA granule often tends to be more peripheral to the Cnn flares of the mother centrosome [17]. Understanding the mechanisms underlying these differences, and testing their influence with respect to centrosome activity, is a promising area of future research.

## Materials and methods

### Fly stocks

The following stocks and transgenic lines were used: *y^1^w^1118^*(Bloomington *Drosophila* Stock Center (BDSC) #1495) was used as the WT control. Null *plp* mutant germline clones were generated by the FLP/ovoD method using *FRT2A, plp^2172^* recombinant chromosomes [66, 67]. *Dhc^LOA^* is a hypomorphic mutation in the dynein heavy chain (*Dhc64C)* gene defined by an F597Y mutation within Dhc (modeled after the murine *Dync1h1* F580Y mutation, legs-at-odd-angles (LOA) [68]. *Ubi-GFP-*y*-Tub23C* expresses GFP-y Tub under the *Ubiquitin* (*Ubi*) promotor [26]; *Ubi-Asl-YFP* expresses *Asl-YFP* under the *Ubi* promoter [69]; *P_BAC_-GFP-Cnn*, expresses Cnn tagged at the N-terminus with EGFP under endogenous regulatory elements [24]; *mCherry-Cnn* expresses Cnn tagged with mCherry with endogenous regulatory elements [70]; *Plp-GFP* is an in-frame C-terminal GFP knock-in at the Plp endogenous locus generated via CRISPR [57]. *UAS-PLP^FL^-GFP* (*Plp^FL^-GFP*) expresses full-length PLP isoform PF under the control of upstream activating sequence (*UAS*) sites [58]; the truncated *Plp* lines, including ΔF1, ΔF2, ΔF5, all express truncated Plp isoform PF fragments under the UAS promoter and are C-terminally tagged with GFP [58]. The *maternal* α*-Tub GAL4* (mat-GAL4; BDSC #7063) driver was used to drive the expression of all UAS transgenes. To examine maternal effects, mutant or transgenic embryos are progeny derived from mutant or transgenic mothers. Flies were raised on Bloomington formula ‘Fly Food B’ (Lab-Express; Ann Arbor, MI), and crosses were maintained at 25°C in a light and temperature-controlled incubator chamber.

### smFISH detection

Stellaris *Plp* and *GFP* smFISH probes conjugated to Quasar 570 or 670 dyes (LGC Biosearch Technologies; Middlesex, UK) were designed against the coding region for each gene using the Stellaris RNA FISH probe designer [17, 71, 18]. smFISH probes were dissolved in nuclease-free water at 25 μM and stored at -20°C before use.

smFISH experiments were performed as previously described using RNase-free solutions [17, 71, 18]. Fixed embryos were rehydrated and washed first in 0.1% PBST (PBS plus 0.1% Tween-20), then in wash buffer (WB; 10% formamide and 2× SSC supplemented fresh each experiment with 0.1% Tween-20 and 2 μg/mL nuclease-free BSA)., then incubated with 100 μL of hybridization buffer (HB; 100 mg/mL dextran sulfate and 10% formamide in 2× SSC supplemented fresh each experiment with 0.1% Tween-20, 2 μg/mL nuclease-free BSA, and 10 mM ribonucleoside vanadyl complex (RVC; S1402S; New England Biolabs; Ipswich, MA) for 10–20 min in a 37°C water bath. Embryos were then incubated in 25 μL of HB containing 0.5 μM smFISH probes in a 37°C water bath overnight. Probes used in this study are listed in Table S2. Embryos were washed three times for 30 min in prewarmed WB, stained with DAPI (1:1000) for 1 hr at room temperature, washed with 0.1% PBST, and mounted with Vectashield mounting medium (H-1000; Vector Laboratories; Burlingame, CA). Slides were stored at 4°C and imaged within 1 week.

### Dual smFISH and immunofluorescence

Dual smFISH and IF experiments were optimized for maintaining the integrity of RNA signals, as previously described [17, 18]. All the following steps were performed with RNase-free solutions. Fixed embryos were processed exactly as described above for smFISH, except for the addition of primary antibody at the same time embryos were incubated overnight in 25 μL of HB containing 0.5 μM smFISH probes in a 37°C water bath. On the next day, embryos were washed four times for 30 min in prewarmed WB, stained with secondary antibody and DAPI (1:1000) for 2 hr at room temperature, washed with 0.1% PBST, and mounted with Vectashield mounting medium (H-1000; Vector Laboratories). Slides were stored at 4°C and imaged within 1 week.

### Microtubule depolymerization and recovery assay

0.5-2.5 hr *YFP-Asl* embryos were collected and dechorionated with bleach for 30 s. The dechorionated embryos were incubated on ice for 5 min to disrupt the microtubules. Embryos were then either immediately fixed in a 1:1 solution of heptane:37% formaldehyde for 3 min, or, to permit microtubule regrowth (recovery), embryos were incubated in room-temperature PBS for 5 min before the fixation. After fixation, all embryos were rinsed in PBS and manually devitellinized as described [17].

### Translational inhibition

To inhibit translation, embryos were treated with inhibitors diluted in Robb’s medium (1 mM calcium chloride, 10 mM glucose, 100 mM HEPES (pH 7.2), 1.2 mM magnesium chloride, 55 mM potassium acetate, 40 mM sodium acetate, and 100 mM sucrose) [72]. To begin, 0.5–2.5 hr embryos were collected and incubated in a 1:1 solution (450 µl: 450 µl) of heptane: Robb’s medium with the appropriate drug or an equivalent volume of DMSO [22]. The concentrations and duration of treatment for each drug are: 3 mM puromycin (Sigma-Aldrich P8833) for 10 min; 0.1 mM anisomycin (Sigma-Aldrich A9789) for 15 min; 0.71 mM cycloheximide (VWR, 97064-724) for 15 min. After drug incubation, Robb’s medium was removed, and 450 µl of 4% paraformaldehyde in PBS was added, and embryos were fixed for 20 min before devitellinization.

### Immunofluorescence

For immunofluorescence with Asl and Cnn antibodies, embryos were fixed in a 1:1 solution of anhydrous methanol (Sigma, #322415): heptane for 15 s and devitellinized in methanol by shaking. For visualization of MTs, embryos were prepared as previously described [73]. Briefly, embryos were fixed in a 1:1 mixture of 37% paraformaldehyde: heptane for 3 min, rinsed in PBS, and manually devitellinized using 30G PrecisionGlide needles (BD). Fixed embryos were rehydrated, blocked in BBT buffer (PBS supplemented with 0.1% Tween-20 and 0.1% BSA), and incubated overnight at 4°C with primary antibodies diluted in BBT. After washing, embryos were further blocked in BBT supplemented with 2% normal goat serum and incubated for 2 hr at room temperature with secondary antibodies and DAPI (10 ng/ml, ThermoFisher). Embryos were mounted in Aqua-Poly/Mount (Polysciences, Inc.) prior to imaging.

The following primary antibodies were used: guinea pig anti-Asl (1:4000, gift from G. Rogers, University of Arizona), rabbit anti-Cnn (1:4000, gift from T. Megraw, Florida State University), mouse anti-α-Tubulin DM1α (1:500, Sigma-Aldrich T6199). Secondary antibodies: Alexa Fluor 488, 568, or 647 (1:500, Molecular Probes), and DAPI (10 ng/ml, ThermoFisher).

### Microscopy and image analysis

Images were acquired on a Nikon Ti-E system fitted with a Yokagawa CSU-X1 spinning disk head, Hamamatsu Orca Flash 4.0 v2 digital complementary metal oxide-semiconductor (CMOS) camera, Perfect Focus system (Nikon), Nikon LU-N4 solid state laser launch (15 mW 405, 488, 561, and 647 nm) using a Nikon 100x, 1.49 NA Apo TIRF oil immersion objective. The microscope was powered through Nikon Elements AR software on a 64-bit HP Z440 workstation.

Images in Figure 4A and 4E were acquired on a Zeiss LSM 880 AiryScan microscope with a 63x 1.4 NA oil objective (“SR” mode; 2x averaging; 1.32 μs pixel dwell). Raw images were processed with Airyscan joint deconvolution in Zen Blue with varying iterations per channel (15 iterations for *Plp* or *Cen* mRNA, 15 iterations for Cnn, 20 iterations for Asl).

smFISH signals were detected and single molecule normalization was performed as described [17, 71, 18]. Briefly, single-channel .tif raw images were segmented in three dimensions using Python scripts adapted from the Allen Institute for Cell Science Cell Segmenter [74]. Each segmented image was compared with the raw image to validate accurate segmentation. RNA objects of ≥50 pixels in segmented images were identified, and object features were extracted, which included surface coordinates.

Distances were measured from the surface of each RNA object to the surface of the closest centrosome. We calculated the percentage of total RNA within 1 μm from the centriole (Asl) or 0 μm from the PCM (Cnn or γTub) surface and selected 10, 8, 6 and 4 μm as the upper boundary for the pseudo-cell radius for NC 10, NC 11, NC 12, and NC 13; respectively, based on measuring the centrosome-to-centrosome distances from a set of representative images. Later interphase/prophase embryos were selected by their large, round nuclei and separated centrosomes.

Fiji (National Institutes of Health; [75]) was used to rotate, split, or merge channels. Images were cropped and brightness/contrast adjusted using Adobe Photoshop. Figures were assembled in Adobe Illustrator.

### RT-PCR

RNA was extracted from ∼2-5 mg of dechorionated 0–2 hr embryos per biological replicate using TRIzol Reagent (#15596026, ThermoFisher Scientific) and treated with1 μL TURBO Dnase (2 U/μL, # AM1907, ThermoFisher Scientific) prior to RT-PCR. 500 ng of RNA was reverse transcribed to cDNA with the iScript cDNA Synthesis Kit following the manufacturer’s protocol (Bio-Rad, #1708891).

qPCR was performed on a Bio-Rad CFX96 Real-time system with iTaq Universal SYBR Green Supermix (#1725121, Bio-Rad; Hercules, CA). Values were normalized to *RpL32* (*rp49*) expression levels. Ct values from the qPCR results were analyzed and the relative expression levels for each condition were calculated using Microsoft Excel. Three biological replicates and three technical replicates were performed on a single 96-well plate using the following primers:

*rp49* Forward: CATACAGGCCCAAGATCGTG

*rp49* Reverse: ACAGCTTAGCATATCGATCCG

*Plp* Foward: CGCAGCAAGGAGGAGATAAC

*Plp* Reverse: TCAGCCTGCAGTTTGTTCAC

*Cen* Forward: AAAGTACCCCCGGTAACACC

*Cen* Reverse: TGAGGATACGACGCTCTGTG

To detect the relative RNA expression level for *Plp* reporter assays, 50 ng cDNA was amplified by PCR for 30 cycles using Phusion High Fidelity DNA Polymerase (M0530L; New England Biolabs) using the following primers:

*Plp* Forward: CACAAACAGCTCGATCAGGA;

*Plp* Reverse: TCATTTTGAGCAACCAGCAG;

*GFP* Forward: ACGTAAACGGCCACAAGTTC;

*GFP* Reverse: AAGTCGTGCTGCTTCATGTG;

*gapdh* Forward: CACCCATTCGTCTGTGTTCG;

*gapdh* Reverse: CAACAGTGATTCCCGACCAG

### Statistical methods

Data were plotted and statistical analysis was performed using GraphPad Prism (v. 9) software. To calculate significance, distribution normality was first confirmed with a D’Agnostino and Pearson normality test. Data were then analyzed by unpaired t-test, one-way ANOVA test, or the appropriate non-parametric test and are displayed as mean ± SD. Data shown are representative results from at least two independent experiments.

### Protein-protein Complex Prediction

To model the interaction between these Plp and Cnn, we ran AlphaFold2 (2.3.2) using the multimer model on the COSMIC^2^ cloud platform with the amino acid sequences of Plp F2: 584-1357 (isoform RF) and Cnn CM2: 1082-1148. AlphaFold2 generated five predicted models. We used PyMOL (version 2.5.7) to visualize and process images of these predicted models. We compared the similarity between 3D protein structures by calculating the Root Mean Square Deviation (RMSD) using the align function in PyMOL by running the command: *align object1, object2*; where object 1 was the CM2 model predicted by AlphaFold, and object 2 was the published 3D crystal structure of CM2 motif (PDB: 5MVW, chain A).

## Competing interest statement

The authors have no competing interests to declare.

## Data Availability Statement

Uncropped gels from Fig. 3 and S2 are available on FigShare: https://figshare.com/s/103951922143448b05d2 https://figshare.com/s/360dfc97047235a2b18a https://figshare.com/s/71f35163efc18e879e7b

## Acknowledgements

For gifts of reagents, we thank Drs. Clemens Cabernard, Timothy Megraw, Nasser Rusan, Greg Rogers, and Simon Bullock. For technical advice and assistance, we are grateful to Dr. Paul Donlin-Asp, Dr. Carolina Eliscovich, Ms. Shuristeen Joubert, Ms. Jordan Goldy, and Mr. Jovan Brockett. We thank Drs. Brian Galletta and Matthew Hannaford for insightful conversations and Dr. Girish Deshpande for constructive feedback on the manuscript.

This work was supported by the Maximizing Investigators’ Research Award (MIRA R35) from the National Institute of General Medical Sciences (NIGMS) of the National Institutes of Health (NIH) R35GM138183 to JRB, National Science Foundation Graduate Research Fellowship (NSF GRFP) DGE-1842190 to MAQ, and NIH grants K99GM143517 to JF and R01GM138544 to DAL.

## Author contributions

JF– formal analysis, funding acquisition, investigation, methodology, project administration, supervision, software, validation, visualization, writing–original draft, and writing–review & editing.

WT– formal analysis, investigation, visualization, writing–review & editing.

MQ– formal analysis, investigation, visualization, writing–review & editing.

JB– supervision, investigation, writing–review & editing.

DAL– conceptualization, funding acquisition, project administration, supervision, writing–original draft, and writing–review & editing.

## Supplemental Data

**Figure S1. Dynein is not essential for *Plp* mRNA localization.** (A) Amino acid alignment of *Drosophila melanogaster (Dmel)* Plp, mouse (*Mmus*) PCNT, and human (*Hsap*) PCNT (Clustal Omega; https://www.ebi.ac.uk/Tools/msa/clustalo/). The amino acid numbers of Plp and PCNT are listed above and fully conserved (*), strongly similar (:), and weakly similar (.) residues are indicated. The dynein light intermediate chain (DLIC) binding motif in human PCNT is noted (blue). (B) Maximum intensity projections of NC 11 embryos of WT and homozygous *Dhc^LOA^* mutants labeled with *Plp* smFISH probes (magenta), Asl antibodies (green), and DAPI (blue). Dashed box regions mark insets. (C) The percentage of *Plp* mRNA localizing within 1 µm from the surface of Asl. Each dot represents a measurement from N=8 WT and N=7 *Dhc^LOA^* NC 11 embryos; see Table S1. Mean ± S.D. are displayed. n.s. not significant by unpaired student t-test. Scale bars: 5 µm (main panels); 2 µm (insets).

**Figure S2. *Plp* mRNA localization requires the *Plp* CDS.** (A) Maximum intensity projections of NC 11 embryos labeled with anti-Asl antibodies (green), *Plp* smFISH probes in WT, or *GFP* smFISH probes in *Plp-GFP* (magenta). Schematic diagrams of labelled RNAs are shown to the left. (B) The percentage of *Plp* mRNA localizing within 1 µm of Asl from N=7 WT and 10 *Plp-GFP* NC 11 embryos. (C) Relative expression level of endogenous *Plp* RNA in 0–2 hr embryos of the indicated genotypes assayed by RT-PCR. (D) Maximum intensity projections of NC 11 embryos labeled with anti-Cnn antibodies (green), *GFP* smFISH probes (magenta) and DAPI (blue) in the following genotypes: (i) *UAS-Plp^5’UTR^-GFP-Plp^3’UTR^*, (ii) *UAS-GFP-Plp^3’UTR^*, (iii) *UAS-GFP, and* (iv) *UAS-Plp^FL^-GFP*. Transgenes in (ii–v) were expressed using *matGAL4* in the presence of endogenous *Plp*. Insets are enlarged in the upper-right corners. Arrowheads mark *Plp* mRNA enriched at centrosomes. Schematic diagrams of GFP-tagged constructs are shown on the left. (E) Relative expression level of the GFP-tagged reporter RNAs in 0–2 hr embryos of the indicated genotypes was assayed by RT-PCR. Uncropped gels are available at < https://figshare.com/s/360dfc97047235a2b18a and https://figshare.com/s/71f35163efc18e879e7b >. Scale bars: 5 µm (main panels); 2 µm (insets).

**Figure S3. The PCM scaffold permits mRNA localization in early embryos.** Maximum intensity projections of NC 11 control and *cnn^B4^* embryos expressing *Asl-YFP* and labeled with (A) *Plp* or (C) *Cen* smFISH probes (magenta) and DAPI (blue). (B) The percentage of *Plp* mRNA localizing within 1 µm from the Asl surface from N=11 WT and 9 *cnn^B^* NC 11 embryos. The percentage of *Cen* mRNA (D) localizing and (D’) residing within granules (defined as ≥ 4 RNA molecules per granule) within 1 µm from the Asl surface from N=11 WT and 10 *cnn^B4^* NC 11 embryos. (E) The AlphaFold Cnn CM2 predicted structure (gray) was superimposed on the 3D crystal structure of Cnn CM2 (PDB: 5MVW; green) [61]. RMSD = 1.4111, (433 to 433 atoms) out of 490 atoms. (F) Side view and top view images of the top 3-ranked AlphaFold models of the Plp F2–Cnn CM2 interaction. Shown are Plp amino acids 1177-1306 (yellow) and Cnn CM2 (gray). Mean ± S.D. is displayed. ****p<0.0001 by unpaired student t-test. Scale bars: 5 µm (main panels); 2 µm (insets).

**Table S1. List of objects quantified in the figures.** Tabulation of genotypes, embryos, centrosomes, and RNA objects quantified in this study.

**Table S2. smFISH probe sequences.** List of *Plp*, *Cen*, and *EGFP* mRNA probes used in this study.

## Notes

### Competing Interest Statement

The authors have declared no competing interest.

