## Supplementary material for "The PCM scaffold enables RNA localization to centrosomes": Figure S1

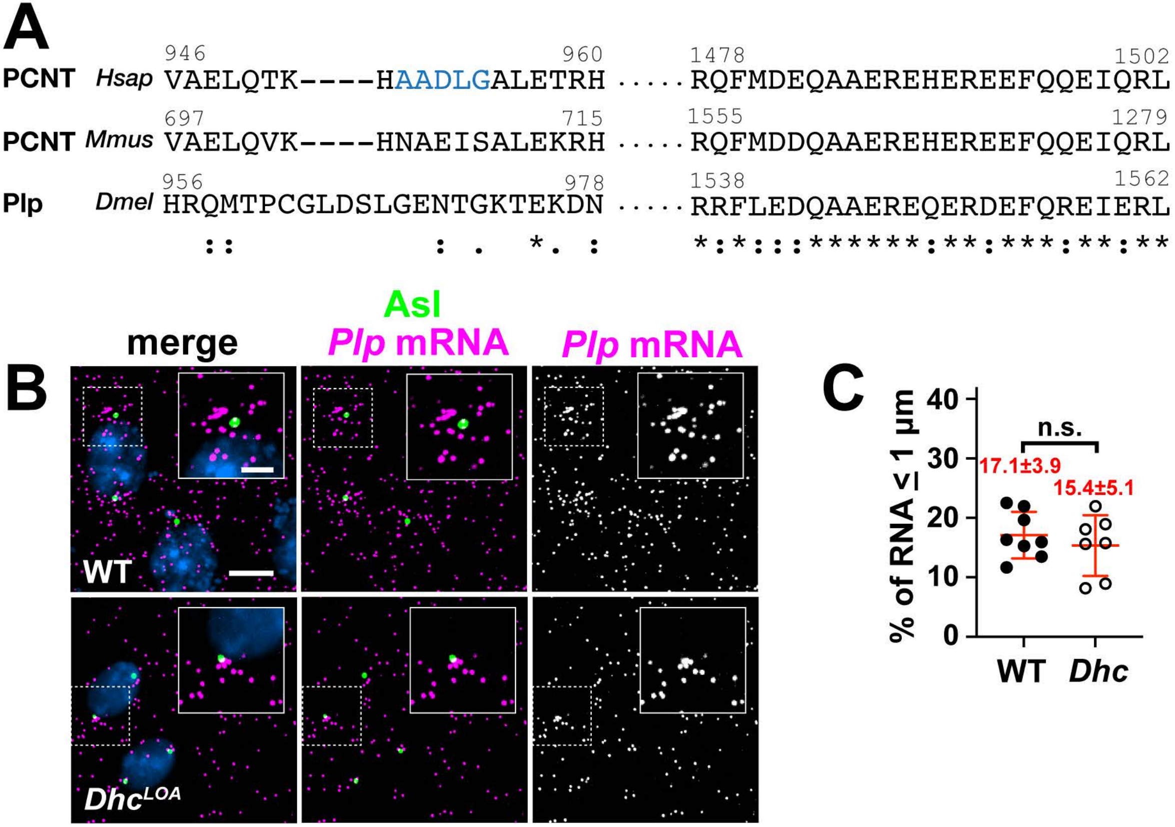


**Figure S1. Dynein is dispensable for *Plp* mRNA localization.** (A) Amino acid alignment of *Drosophila melanogaster* (*Dmel*) Plp, mouse (*Mmus*) Pcnt, and human (*Hsap*) PCNT (Clustal Omega; https://www.ebi.ac.uk/Tools/msa/clustalo/). The amino acid numbers of Plp and PCNT are listed above and fully conserved (*), strongly similar (:), and weakly similar (.) residues are indicated. The dynein light intermediate chain (LIC) binding motif in human PCNT is noted (blue). (B) Maximum intensity projections of NC 11 embryos of WT and homozygous *Dhc*$^{LOA}$ mutants labeled with *Plp* smFISH probes (magenta), Asl antibodies (green), and DAPI (blue). Dashed box regions mark insets. (C) The percentage of *Plp* mRNA localizing within 1 µm from the surface of Asl. Each dot represents a measurement from a single embryo; see Table S1 for N embryo and RNA objects examined. Mean ± S.D. are displayed. n.s. not significant by unpaired student t-test. Scale bars: 5 µm (main panels); 2 µm (insets).
