## Supplementary material for "The PCM scaffold enables RNA localization to centrosomes": Figure S2

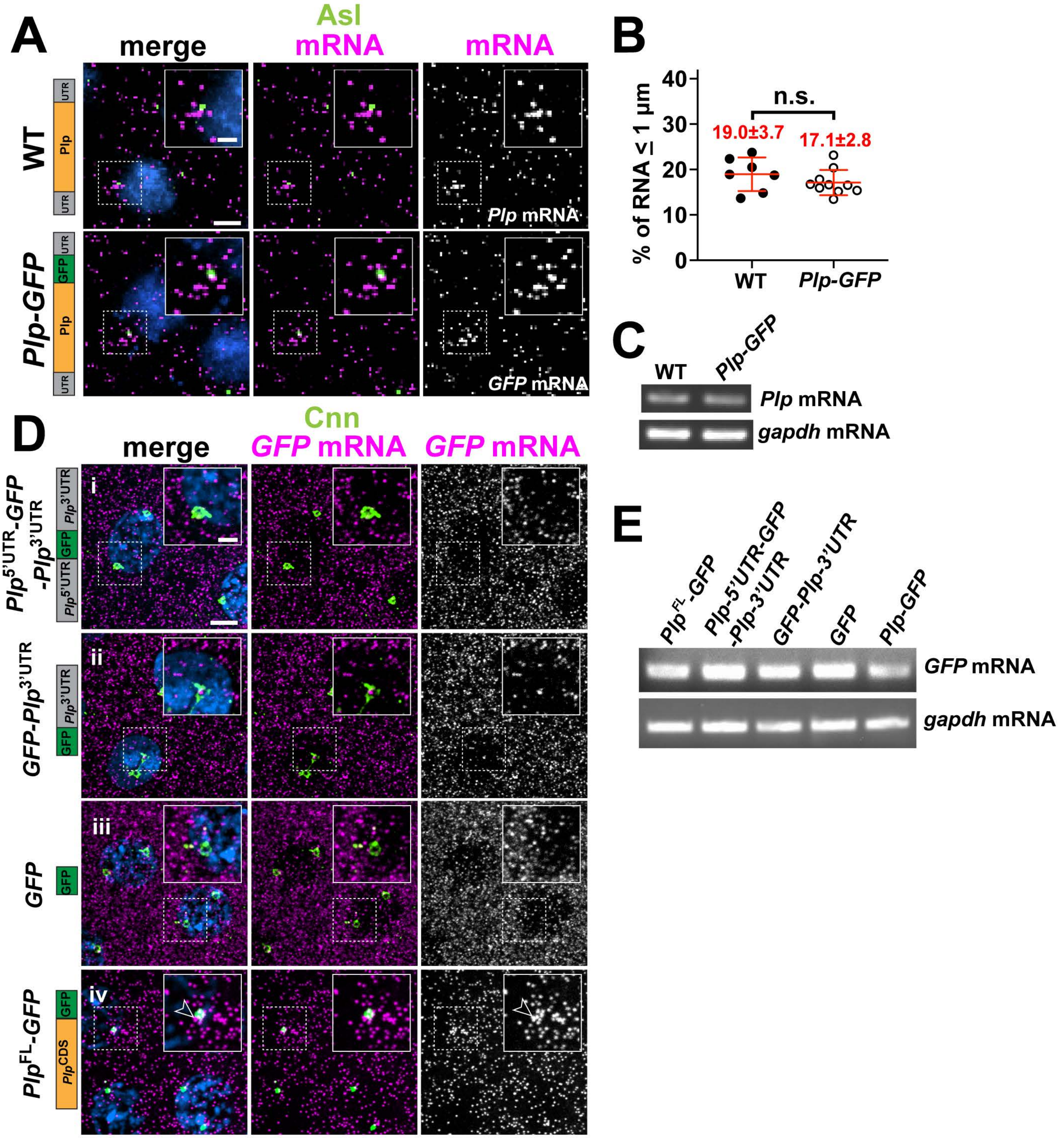


**Figure S2.** ***Plp*** **mRNA localization requires the** ***Plp*** **CDS.** (A) Maximum intensity projections of NC 11 embryos labeled with anti-Asl antibodies (green), *Plp* smFISH probes in WT, or *GFP* smFISH probes in *Plp-GFP* (magenta). Schematic diagrams of labelled RNAs are shown to the left. (B) The percentage of *Plp* mRNA localizing within 1 μm of Asl. (C) Relative expression level of endogenous *Plp* RNA in 0-2 hr embryos of the indicated genotypes as assayed by RT-PCR. (D) Maximum intensity projections of NC 11 embryos labeled with anti-Cnn antibodies (green), *GFP* smFISH probes (magenta) and DAPI (blue) in the following genotypes: (i) *UAS-Plp5'UTR-GFP-Plp3'UTR*, (ii) *UAS-GFP-Plp3'UTR*, (iii) *UAS-GFP*, and (iv) *UAS-PlpFL-GFP*. Transgenes in (ii–v) were expressed using *matGAL4* in the presence of endogenous *Plp*. Insets are enlarged in the upper-right corners. Arrowheads mark *Plp* mRNA enriched at centrosomes. Schematic diagrams of GFP-tagged constructs are shown on the left. (E) Relative expression level of the GFP-tagged reporter RNAs in 0-2 hr embryos of the indicated genotypes was assayed by RT-PCR. Uncropped gels are available at < https://figshare.com/s/360dfc97047235a2b18a and https://figshare.com/s/71f35163efc18e879e7b >. Scale bars: 5 μm (main panels); 2 μm (insets).
