## Supplementary material for "The PCM scaffold enables RNA localization to centrosomes": Figure S3

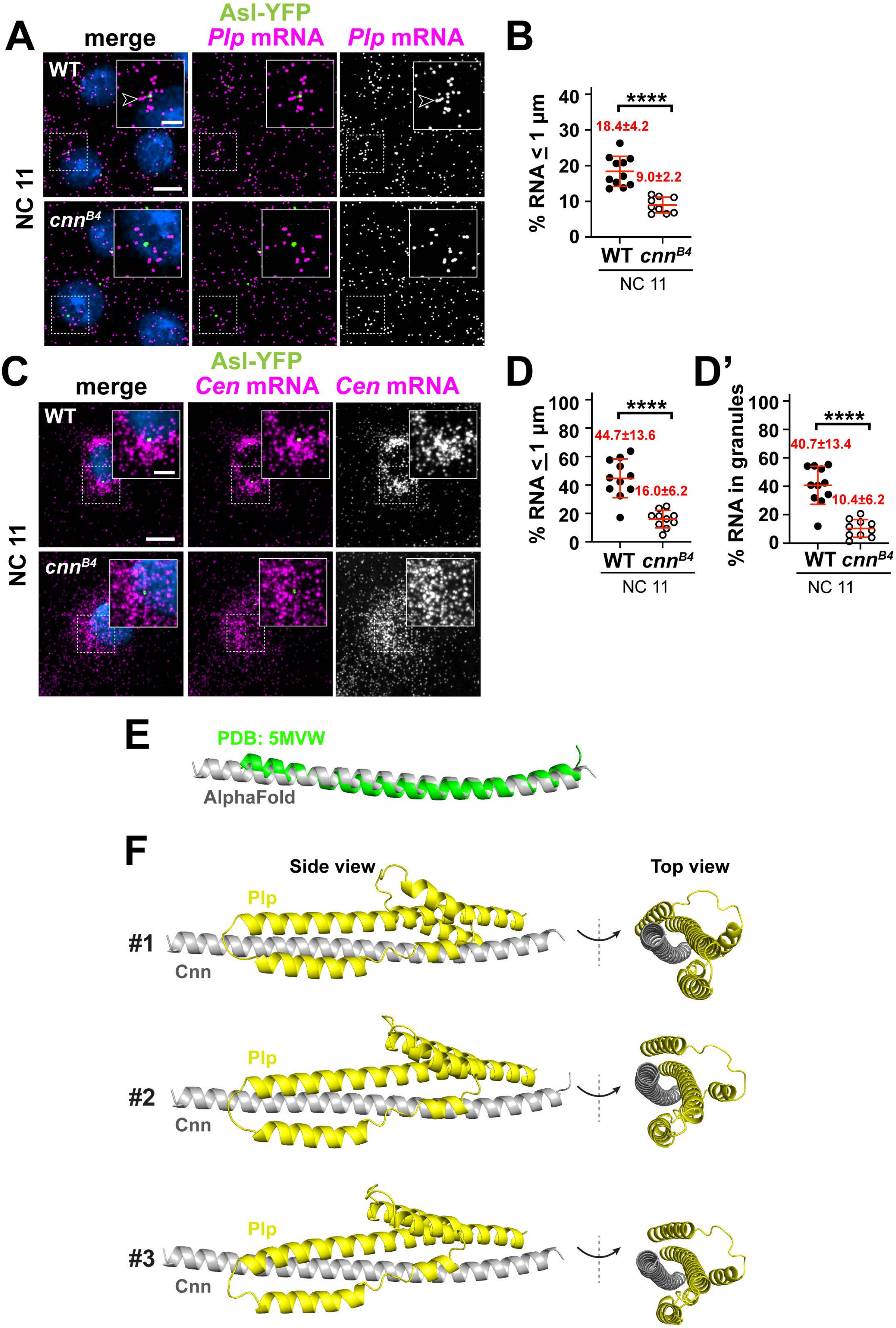


**Figure S3. The PCM scaffold permits mRNA localization in early embryos.** Maximum intensity projections of NC 11 control and *cnn*$^{B4}$ embryos expressing *Asl-YFP* and labeled with (A) *Plp* or(C) *Cen* smFISH probes (magenta) and DAPI (blue). (B) The percentage of *Plp* mRNA localizing within 1 µm from the Asl surface. The percentage of *Cen* mRNA (D) localizing and (D') residing within granules (defined as ≥ 4 RNA molecules per granule) within 1 µm from the Asl surface. (E) The AlphaFold Cnn CM2 predicted structure (gray) was superimposed on the 3D crystal structure of Cnn CM2 (PDB: 5MVW; green) [61]. RMSD = 1.4111, (433 to 433 atoms) out of 490 atoms. (F) Side view and top view images of the top 3-ranked AlphaFold models of the Plp F2–Cnn CM2 interaction. Shown are Plp amino acids 1177-1306 (yellow) and Cnn CM2 (gray). Mean ± S.D. is displayed. ****p<0.0001 by unpaired student t-test. Scale bars: 5 µm (main panels); 2 µm (insets).
